# Conformational flexibility Of A Highly Conserved Helix Controls Cryptic Pocket Formation In FtsZ

**DOI:** 10.1101/2020.09.08.288134

**Authors:** Aisha Alnami, Raymond S. Norton, Helena Perez Pena, Shozeb Haider, Frank Kozielski

**Author notes:** Correspondence and requests for materials should be addressed to F.K.

## Abstract

*Mycobacterium tuberculosis* is responsible for more than 1.6 million deaths per year. Overcoming failure from established therapies owing to multidrug resistance requires the identification of novel targets. One potential antibacterial target is filamentous temperature sensitive protein Z (FtsZ), which is the bacterial homologue of mammalian tubulin, a validated cancer target. *M. tuberculosis* FtsZ function is essential, with its inhibition leading to arrest of cell division, elongation of the bacterial cell and eventual cell death. However, the development of potent inhibitors against FtsZ has been a challenge due to the lack of structural information. Here we have solved multiple crystal structures of *M. tuberculosis* FtsZ in complex with coumarin analogues. Coumarins bind exclusively to two novel cryptic pockets in nucleotide-free FtsZ but not to the binary FtsZ-GTP or GDP complexes. Our findings provide a detailed understanding of the molecular basis for cryptic pocket formation, controlled by the conformational flexibility of the H7 helix, and thus reveal an important structural and mechanistic rationale for coumarin’s antibacterial activity.

---

Tuberculosis (TB) is one of the leading causes of morbidity and mortality caused by an infectious agent. The World Health Organization estimates that the latent form of TB has infected nearly one-quarter of the world’s population. Ten million people have developed the disease (equivalent to 142 cases per 100,000 population), and about 1.3 million died in 2017. Additionally, 558,000 reported cases were classified as multidrug resistant tuberculosis (MDR-TB) patients, and 8.5% of these patients developed extensively drug-resistant tuberculosis (XDR-TB)[1].

After a 40-year gap, the US Food and Drug Administration (FDA) approved two new anti-TB agents, Bedaquiline in 2012 [2] and Delamanid in 2014 [3], to be used as adjuncts to current anti-TB agents in treating MDR-TB[4]. However, multidrug-resistant TB infections remain a substantial threat to public health worldwide, creating a pressing need to explore novel anti-TB agents with new mechanisms of action.

FtsZ, the bacterial homologue of mammalian tubulin, is highly conserved among all types of bacteria, including Mycobacterium strains [5]. FtsZ is an essential cell division protein, with both GTPase and polymerisation activities. In a GTP-regulated manner, FtsZ subunits polymerise at the centre of the cell into protofilaments to form a dynamic ring-like structure called the Z-ring [6]. The Z-ring functions as a scaffold for the assembly of the divisome, a multiprotein complex that causes contraction of the Z-ring, finally resulting in septum formation and cell division [7, 8]. Abnormalities in polymerization and/or GTPase activity result in inhibition of the Z-ring and septum formation, leading to inhibition of cell division and cell death.

In the effort to develop inhibitors of bacterial cell division that target FtsZ, several groups have identified inhibitors based on natural products, synthetic small molecules, and nucleotides. However, many of these hits have been described as “false positives” or possessing “irreproducible activity” [9, 10], and progress of the majority of these projects has been hampered by the lack of available structural data such as crystallographic FtsZ-inhibitor complexes for structure-based drug design. It has been proposed that it may be an “intrinsic challenge” to crystallise FtsZ in complex with inhibitors [11]. Hitherto, only a single binding site for PC190723 and related compounds [12] has been identified in *Staphylococcus aureus* FtsZ, but not in any other bacterial species [13].

In this study we successfully identified two novel inhibitor-binding pockets in FtsZ from *M. tuberculosis* in crystal structures with a resolution of 1.7 Å. We show that these pockets are cryptic in nature since they appear in nucleotide-free FtsZ but are absent in the nucleotide-bound form. Furthermore, their formation and dissolution depend on the structural dynamics of H7, a highly conserved helix across all bacterial FtsZ.

## Results

### Overall description of the binary FtsZ-nucleotide bound structure

We determined the structure of *M. tuberculosis* FtsZ in the absence and presence of the coumarin analogue 4-hydroxycoumarin (Fig. 1a). Data collection and refinement statistics are summarised in Tables S1 and S2. FtsZ crystallises as a dimer, with one binary FtsZ-nucleotide complex and one molecule of nucleotide-free FtsZ. Since we did not add GTP or GDP to the purification buffer or the crystallisation conditions the nucleotide originates from expression in *E. coli*. FtsZ contains an N-terminal enzymatic domain and a shorter C-terminal domain designated the GTPase-activating domain, which are separated by a long helix named H7 (Fig. 1b) that is structurally conserved across all bacterial species (Fig. S1). The nucleotide-binding pocket is located in the N-terminal domain (helices H1-H6 and strands β1-β6), while the C-terminal domain (H8-H10 and β7-β10) contains the important T7 loop at the tip of helix H7, responsible for the binding of various auxiliary proteins (Fig. 1b).

**Fig 1.**
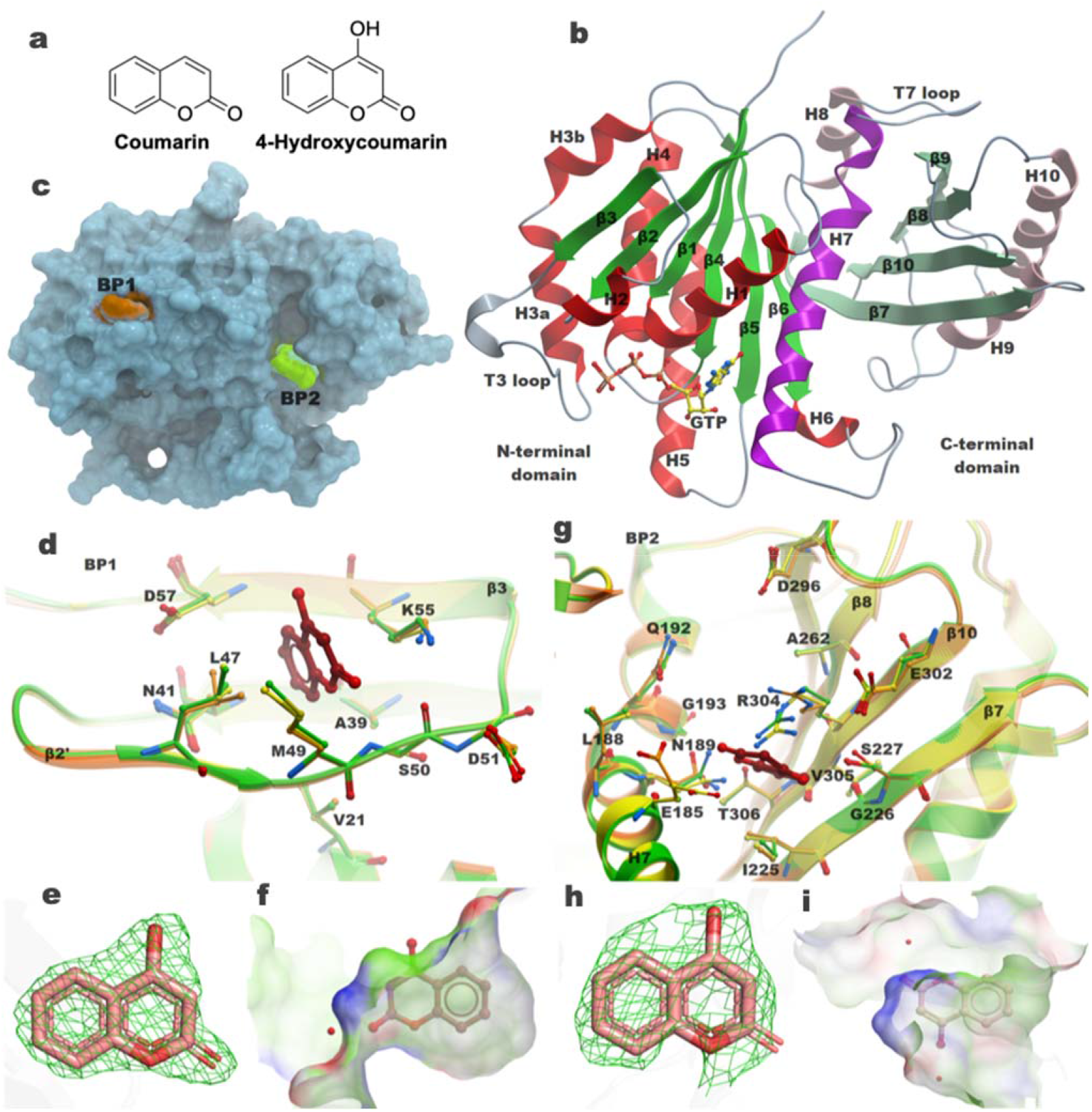
Structures of FtsZ-inhibitor complexes. **a**, Chemical structures of coumarin and 4-hydroxycoumarin as known inhibitors of the bacterial cell division protein FtsZ. **b**, Overall structure representing the binary GTP-bound form of *M. tuberculosis* FtsZ. The N-terminal enzymatic domain is coloured in red (helices) and green (β-strands), whereas the C-terminal GTPase-activating domain is coloured in pink and light green for helices and β-strands, respectively. Loop regions in both domains are shown in grey. The central helix H7 separating the two domains is coloured in purple. The bound nucleotide GTP is shown as a liquorice model. **c**, Surface representation of Nucleotide-free FtsZ highlighting the position of binding pockets (BP) 1 and 2. **d**, Interactions of 4-hydroxycoumarin in BP1. Residues surrounding the inhibitor are illustrated in stick representation and 4-hydroxycoumarin is coloured in red. Hydrogen-bond interactions are displayed as dotted lines. **e**, Difference electron density map of the inhibitor contoured at 1σ fully covering the inhibitor in BP1. **f**, Volume of BP1 with bound 4-hydroxycoumarin and a water molecule **g**, Molecular interactions of 4-hydroxycoumarin in BP2. Residues surrounding the inhibitor and 4-hydroxycoumarin is coloured in red. Hydrogen-bond interactions are displayed as dotted lines. **h**, Electron density of the inhibitor contoured at 1σ fully covering the inhibitor in BP2. (**i)** Volume of BP2 with bound 4-hydroxycoumarin and two water molecules.

### Molecular interactions of 4-hydroxycoumarin with FtsZ

4-hydroxycoumarin binds exclusively to nucleotide-free FtsZ, and not to the nucleotide-FtsZ complex. It binds to two novel inhibitor-binding pockets not previously identified in FtsZ (Fig. 1c). Binding pocket 1 (BP1) is formed by secondary structure elements β2, β2’ and β3 (Fig. 1d). 4-hydroxycoumarin binds with full occupancy and the electron density fully covers the entire molecule (Fig. 1e; Fig. S2). The benzyl ring is oriented inwards, towards the more hydrophobic part of the inhibitor-binding pocket. The 4-hydroxy group points towards the solvent and does not form any interactions with FtsZ, although it is in close proximity to several water molecules. (Fig. S3). The benzyl group forms hydrophobic interactions with residues Leu47 of β2’ and Met49, from the turn between β2’ and β3 and Lys55 and Asp57 of β3, with 4-hydroxycoumarin aligned almost parallel to the elongated side chain of Lys55. 4-hydroxycoumarin forms two hydrogen bond interactions with residues of the inhibitor-binding pocket including the carbonyl oxygen, which interacts with the nitrogen atom of the Lys55 side chain (3.66 Å) and the backbone nitrogen of Ser50 (3.01 Å). A third interaction with the side chain nitrogen of Lys55 is mediated via a structural water molecule. The ring oxygen of the pyrone group also interacts with the main chain nitrogen of Ser50 (3.04 Å). The volume of BP1 is 104.4 Å^3^ (Fig. 1f).

A second 4-hydroxycoumarin binding pocket (BP2) appeared later during refinement. 4-hydroxycoumarin in BP2 is sandwiched between Glu185, Asn189, Gln192 of helix H7 and Ile225 (strand β7), Ala262 (strand β8), Glu302 and Arg304 (strand β10) (Fig. 1g). Similar to BP1, the benzyl group points towards the protein interior, establishing various hydrophobic interactions, whereas the more hydrophilic pyrone ring system points towards the solvent, establishing several water-mediated hydrogen-bond interactions. The electron density around the inhibitor is well defined and 4-hydroxycoumarin binds in the second pocket with full occupancy (Fig. 1h, Fig. S2). There are several short- and long-distance hydrogen-bond interactions between residues of FtsZ and water molecules. Long-distance interactions (3.6 to 4.9 Å) occur between the 4-hydroxy group and the side chain oxygen atoms of Ser227, Glu302 and Arg304, as well as between the pyrone oxygen atoms with main chain carbonyl oxygen and the carboxyl side chain of Glu185. The 4-hydroxy substituent establishes a hydrogen bond interaction with a structural water molecule whereas the two oxygen atom of the pyrone ring show a second interaction with a water molecule (Fig. S3). The volume of BP2 is 146.6 Å^3^ (Fig. 1i).

### Structural changes associated with 4-hydroxycoumarin binding

4-hydroxycoumarin binds in two distinct pockets. We describe these as cryptic pockets because they are present exclusively in the nucleotide-free state but absent in nucleotide-bound FtsZ. So what is the structural rationale for this selectivity of 4-hydroxycoumarin for nucleotide-free FtsZ over the nucleotide-bound form? Overlaying the structure of nucleotide-free FtsZ-coumarin binary complex with that of the FtsZ-nucleotide complex (Fig. 2) reveals significant structural differences. BP1 for 4-hydroxycoumarin in the nucleotide-free structure is formed by strands β2, β2’ and β3, whereas in the FtsZ-nucleotide complex strand β2’ converts into a 2-turn helix H2 (Fig. 2a). This structural rearrangement from a β-strand to a helix leads to a shift of Leu47 by 3.15 Å such that its side chain is projected into BP1, thereby restricting the binding of 4-hydroxycouamrin. Additionally, Ser50, whose main chain nitrogen forms hydrogen bond interaction with the oxygen atom of the pyrone ring is shifted by 2.49 Å. Thus, BP1 for 4-hydroxycoumarin in nucleotide-free FtsZ is absent in the nucleotide-bound form.

**Fig 2.**
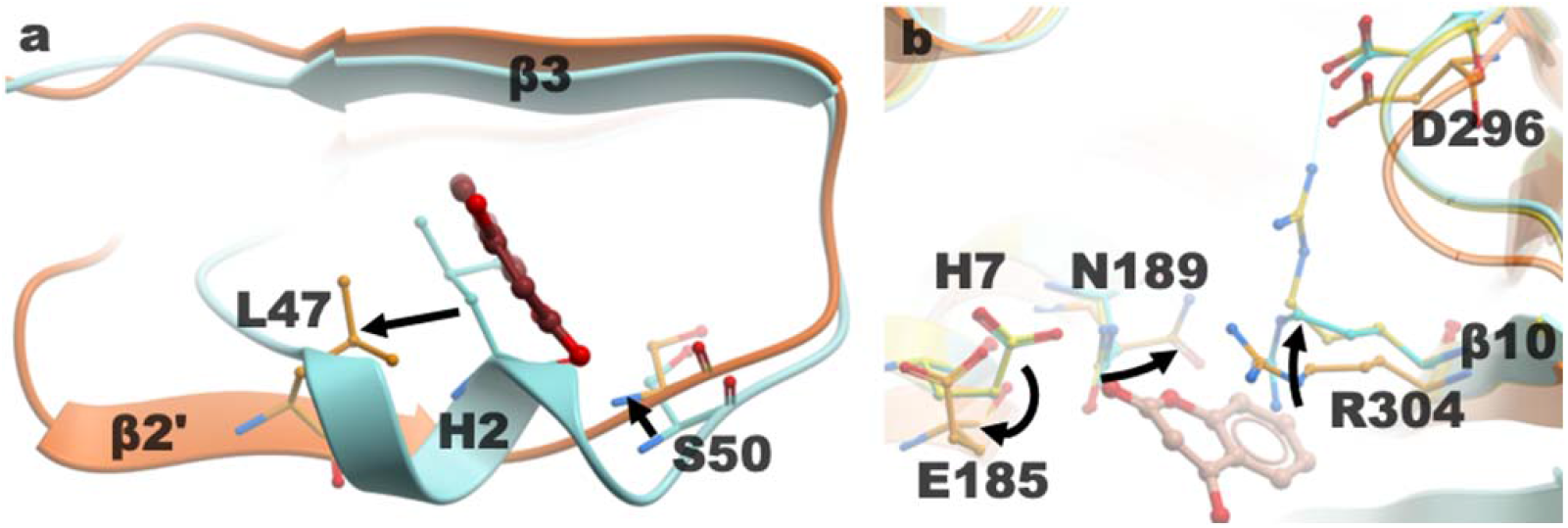
Structural differences of the 4-hydroxycoumarin binding pockets between the nucleotide-bound (cyan/yellow) and the nucleotide-free form of FtsZ (orange), 4-hydroxycoumarin (red) is shown as a liquorice model. **a**, Comparison of the BP1 in nucleotide-free and -bound FtsZ. 4-hydroxycoumarin would sterically clash with helix H2 in the nucleotide-bound form. **b**, Overlay of BP2 in the nucleotide-free form and the nucleotide-bound proteins. Arrows represent the reorientation of the side chains around the binding site, allowing 4-hydroxycoumarin to bind.

When overlaying the nucleotide-free and nucleotide-bound structures to compare BP2 (Fig. 2b), the most obvious difference is the side chain conformation of Arg304. The guanidinium side chain runs parallel to the inhibitor when 4-hydroxycoumarin is bound, but it can reorient itself to form a new hydrogen bond interaction with Asp296 or towards β7 and thus occlude BP2. Also, in the nucleotide-bound form, the side chains of Glu185 and Asn189 are in closer proximity to the putative 4-hydroxycoumarin binding pocket and would have to move away to make space for the inhibitor.

### Comparison of the 4-hydroxycoumarin binding pockets

An important question is whether the two new cryptic binding pockets are already present in the native nucleotide-free form, or if the inhibitor induces the formation of the binding pockets? To be able to directly compare the 4-hydroxycoumarin binding pockets, we superimposed nucleotide-free FtsZ and the nucleotide-free FtsZ-coumarin complexes (Fig. 1d,g; Table S1). Both native and coumarin-bound complexes crystallise in the same space group with very similar cell parameters displaying the same crystal contacts. In both nucleotide-free structures, the residues of the BP1 are essentially identical. The only noticeable difference is a slight change in conformation of the Leu47 side chain, which slightly rotates to orient itself more favourable with respect to the 4-hydroxycoumarin (Fig. 1d). The native nucleotide-free structure contains four water molecules bound in place of 4-hydroxycoumarin.

The situation is distinct for BP2 since significant differences can be observed: in the native structure the side chains of Ile225 and Arg304 clearly adopt two alternative conformations where one of those extends into the BP2. Two additional side chains (Glu185 and Asn189) also display conformational changes (Fig. 1g).

### Conformational dynamics of FtsZ states

To provide a structural rationale for the selectivity of 4-hydroxycoumarin towards the nucleotide-free FtsZ over the nucleotide-bound form and understand the differences in the conformational dynamics of the complexes, we carried out long timescale molecular dynamics simulations. Two sets of statistically independent simulations for nucleotide-bound (GTP and GDP) and nucleotide-free states were simulated for 3 μs each. During the simulations, the nucleotides were retained in the complexes in their bound state. However, the 4-hydroxycoumarin was removed from its corresponding binding pockets in the nucleotide-free state. This was done to study the transient nature of the binding pockets and exclude any influence of the ligand on the formation or disappearance of the cryptic pockets in the simulated system. We analysed the dynamic interconversion of structural conformations between distinct FtsZ states and identified relevant intermediates, via RMSD based clustering of all MD simulation trajectories using the k-medoids algorithm (Fig. S4) [14].

Notably, there was no exclusive unidirectional relationship between nucleotide-free and nucleotide-bound states. The final ensembles of the nucleotide-free form could not be reached from the GTP or GDP-bound states directly. BP1 is fully formed when strand β2’ is present (cluster 4), while it is absent in nucleotide-bound states when a complete helix H2 is present (clusters 1 and 6) (Fig. S5). The intermediate clusters highlight varying levels of incomplete helicity of helix H2, multiple conformations of the loop T3 (between β3 and helix H3), which interacts with the phosphate groups in the nucleotide-bound state and the loop between β6 and H7 helix (Fig. S6). A more detailed analysis of the clusters indicated that the long helix H7 (Residues 176-199) is also highly dynamic and prone to helical bending. In the nucleotide-free state, helix H7 is bent towards the C-terminal domain, away from the nucleotide-binding site, where H7 helix did not show considerable kinking in the nucleotide-bound states. This suggested that helix H2, which forms the boundary of BP1, behaved like a constrained spring that was wound when helix H7 was straight (nucleotide-bound states) and relaxed into the β2’-strand when helix H7 moved away towards the C-terminal domain in the nucleotide-free state. The linked inter-conversion of helix H2 and the β2’ strand, concomitant with helical H7, bending resulted in the formation or disappearance of BP1.

### Helix H7 bending controls the formation of cryptic binding pockets

Flexible helices frequently support conformational transitions between different functional states of proteins [15] and tend to be conserved in evolution, implicating them in functional roles. Of all the amino acids, glycine and proline have the strongest destabilising effect on helices [16, 17]. Glycine in particular disfavours helix folding since the entropic cost of tethering glycine in a helix exceeds the gain of enthalpy upon helix formation [18-20]. H7 helical bending was calculated as a function of the nucleotide-bound/free states from the simulations. Analysis was carried out separately on each simulated state.

Helix H7 distortion distribution was then extrapolated as a heat map and was colour coded according to the local angle magnitude (Fig. 3a). The flexibility of helix H7 is greater in the nucleotide-free state, followed by GDP-bound and GTP-bound states. In the GTP-bound state, the C-terminal end of helix H7 is tethered to neighbouring structural elements via Gln192 making hydrogen bond interactions with Gln30 of helix H1 and Arg304 (β10) (Fig. 4a). Mild flexibility is observed between residues 189-195, with an average kink of 12°-18°. Furthermore, the N-terminal region of helix H7 forms part of the nucleotide-binding site. The side chain of Asp184 establishes hydrogen bond interactions with the guanine nucleotide, while Phe180 makes a π-stacking interaction with the imidazole ring of the purine moiety. There are a number of other hydrogen bond interactions, for example between the γ-phosphate and Thr42 (loop between β2 and helix H2) and the backbone nitrogen atom of Ala68 and Ala70 (loop between β3 and helix H3), which constrain the nucleotide within its binding site and indirectly restrict helix H7 bending. It is also noteworthy that Thr42 sits at the edge of helix H2, which in the nucleotide-bound form prevents the formation of BP1.

**Fig 3.**
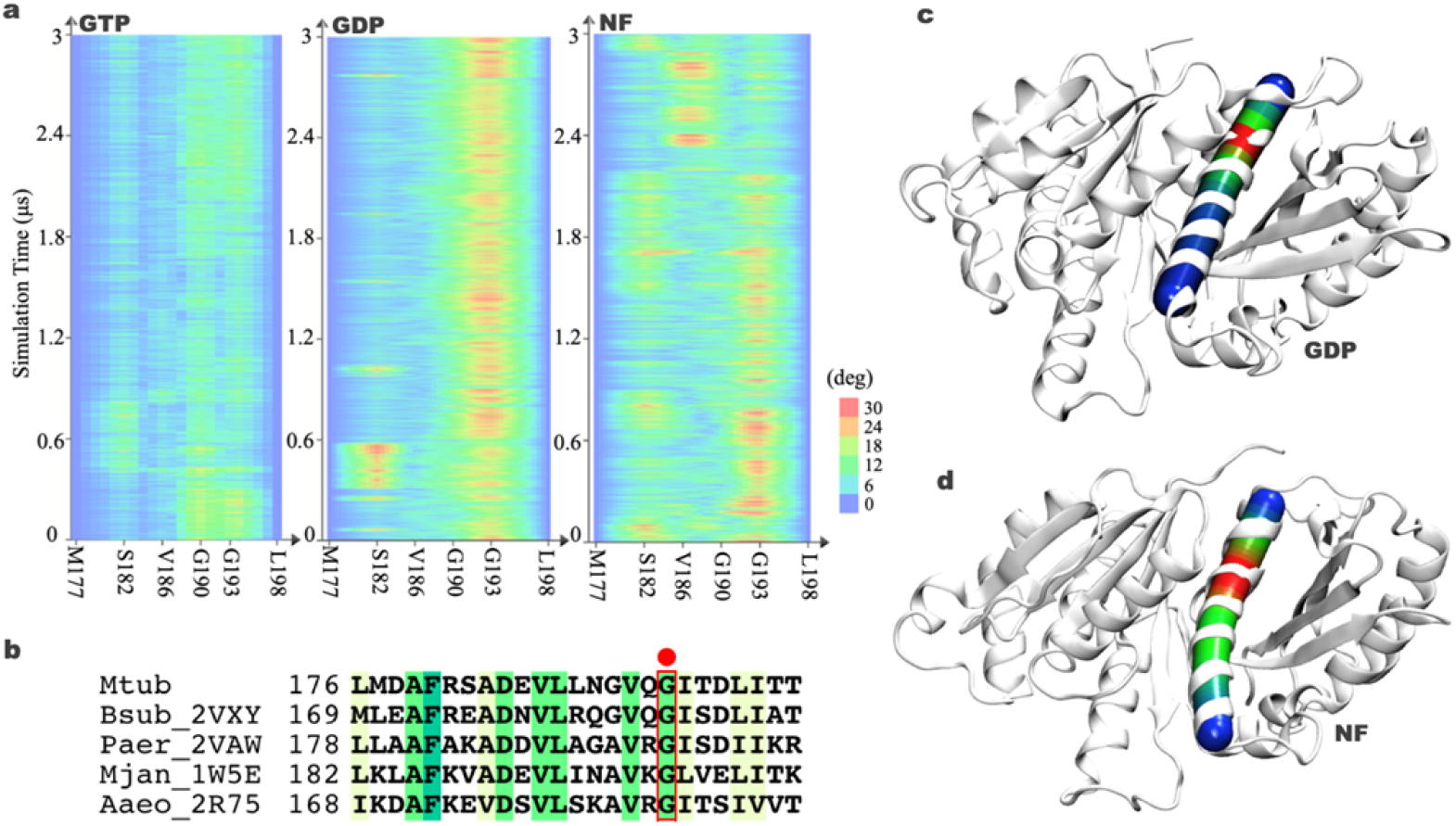
Dynamics of helical H7 bending. **a**, Heat map of H7 helix bending in the FtsZ-GTP (left), FtsZ-GDP (middle) and nucleotide-free FtsZ (right) states plotted as a function of simulation time. **b**, Structural conservation of helix H7 highlighting the position of Gly193 (red box) in *M. tuberculosis* (current structures), *B. subtilis* (PDB id 2VXY), *P. aeruginosa* (PDB id 2VAW), *M. jannaschii* (PDB entry 1W5E) and *A. aeolicus* (PDB entry 2R75). Conformational snapshot at 3 μs of **c** FtsZ-GDP and **d** the nucleotide-free complex illustrating helical bending. Helix H7 is represented as a cylinder. The colour legend representing the kink is the same as in Fig. 3a.

**Fig 4.**
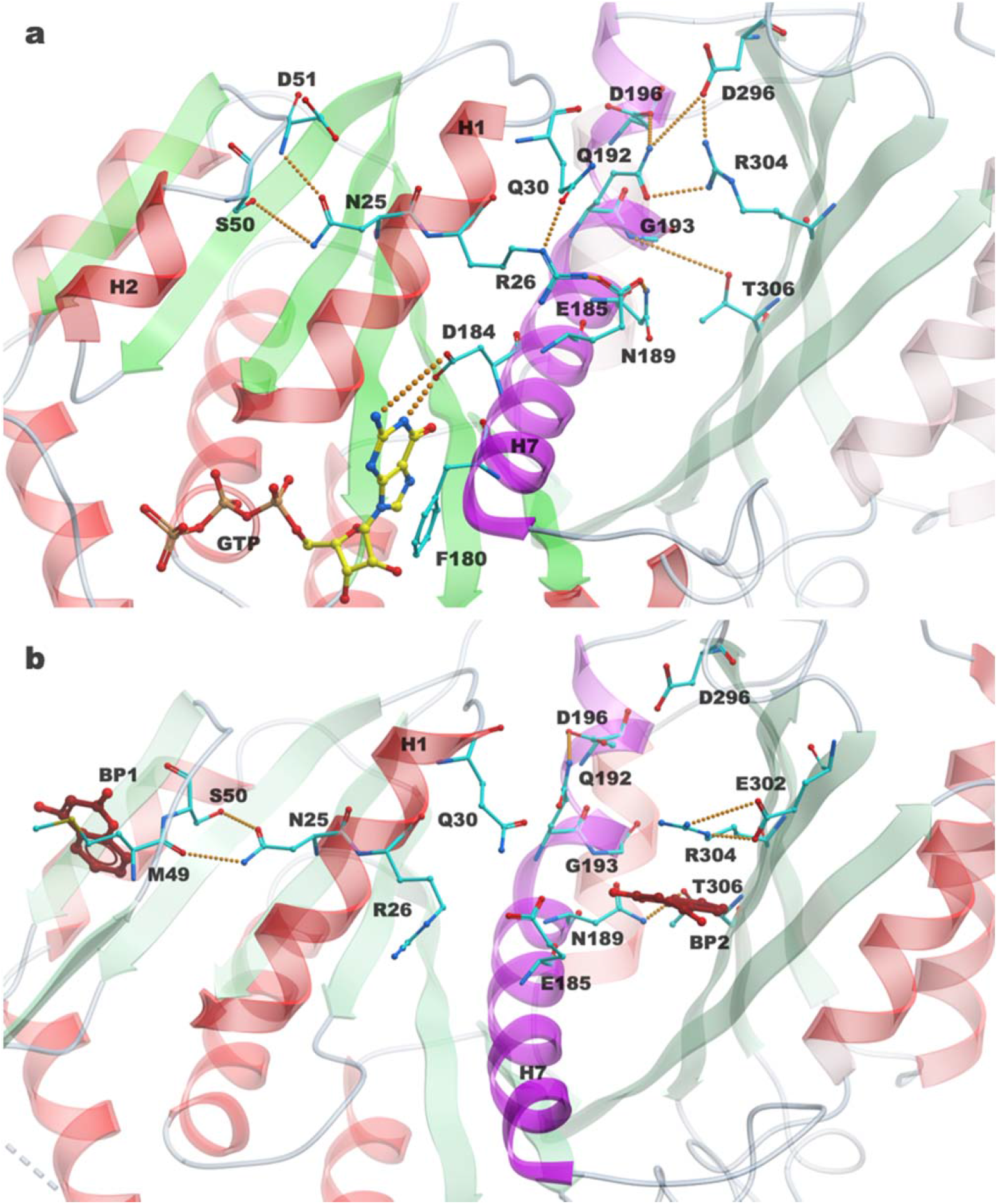
Tethering of BP1 and BP2 via H7 helix in **a** nucleotide-bound and **b** nucleotide-free FtsZ structures. Helix H7 is coloured purple, GTP is coloured yellow and 4-hydroxycoumarin are coloured red. Orange spheres represent hydrogen bond interactions.

In the GDP-bound state, we observe similar tethering of helix H7 with H1, although interactions with β10 are not present. Helical bending is observed between residues 190-194 and is most pronounced at Gly193 (Fig. 3a). During sampling of the first 600 ns, additional helical bending is also observed around Ser182. The loss of a γ-phosphate in GDP results in loss of the interaction with Thr42 and Ala68 and the additional volume generated permits enhanced flexibility of helix H7. The greatest variation in helical flexibility is seen in the nucleotide-free state. In addition to Ser182 and Gly193, a helical kink is also seen at Val186. Comparing helix H7 flexibility across the three states indicated that bending occurs predominantly at Gly193, a highly conserved residue in H7 helix across the bacterial FtsZ family (Fig. 3b). This suggests that the helical movement of H7 is conserved and possibly common to all bacterial FtsZ proteins. Gly193 is positioned near the C-terminal end of helix H7 and act as a hinge point for kink movements, thereby exerting its effects on the farther N-terminal part of helix H7, adjacent to the nucleotide-binding site. The nucleotides act like a wedge that restricts H7 helical motion. Therefore, in the nucleotide-free state, G193 acts like a primary pivot point of helix H7, localizing on its concave side (Fig. 3d). A comparison of the different states, from both crystallographic structures and MD simulation trajectories, suggests that helical bending is heavily influenced by steric hindrance of the nucleotide and affects the collective dynamics of helix H7 at its N-terminal apex.

Is the formation of BP2 in the nucleotide-free state also linked with the dynamics of helix H7? In the GTP-bound state, Glu185 makes hydrogen bonds with Asn189, which is stable throughout the course of the simulation. Ile225, whose hydrophobic side chain forms the base of BP2 is positioned opposite Asn189. In addition, the side chain of Arg304, which in the crystal structure points into BP2, makes interactions with Gln192 and Asp296. Gln192 also forms stable hydrogen bond interactions with Gln30. These interactions strongly tether helices H1 and H7 and β9 together. This tethering binds the local structural elements from the N-terminal and C-terminal domains together and prevents the formation of BP2.

In the GDP-bound state, tethering is observed between helices H1 and H7 through residues Arg26 and Glu185, Gln30 and Gln192, Lys33 and Asp196. However, interactions between helix H7 and Arg304 of β10 are lost. Even though the volume of BP2 increases slightly from that observed in the GTP-bound state, it is still not large enough to accommodate 4-hydroxycoumarin.

In contrast, in the nucleotide-free state, the conformational dynamics of helix H7 prevent any stable interactions or tethering of the local structural elements together at its C-terminal end. As a result, the side chains of Arg304 and Asp189 can adopt multiple conformations. The effects of H7 helical bending on its N-terminal region allow the side chain of Glu185 to rearrange and form the boundary of BP2. The multiple side chain reorganisations impart a defined shape to BP2, making it large enough to accommodate 4-hydroxycoumarin. In summary, the formation of BP1 and BP2 are directly linked with the structural dynamics of helix H7 in the nucleotide-free state.

### 4-hydroxycoumarin binding sites are involved in pocket crosstalk

Small molecule binding sites in proteins have been shown to sometimes contribute to protein allosteric signalling [21, 22]. Since both 4-hydroxycoumarin binding pockets appear together in the nucleotide-free crystal structure and the dynamics of helix H7 bending contributes to the formation and dissolution of both BP1 and BP2, we investigated whether there was any communication between the two sites. We used protein conformations extracted from MD trajectories to detect the formation and spatiotemporal evolution of the pockets formed in the nucleotide-free state of FtsZ. Such communication termed as “pocket crosstalk” has been effectively used to study pharmaceutically relevant targets, including the purine nucleoside phosphorylase, adenosine receptor in a membrane environment and the kinase family [23]. A merging and splitting frequency matrix was calculated into a 3D network graph, where nodes represent the pockets and the edges indicate communication between two pockets or how often two pockets exchange atoms. A total of five pockets was identified that existed for more than 24% during the course of the nucleotide-free simulation run. Pocket identifier (pID) 7 (green) corresponds to BP1, pID 4 (blue) is BP2, pID 3 (red) corresponds to the nucleotide binding site and pID 20 (pink) and 14 (orange) are two sub-pockets that form as a result of the fluctuations seen in loop T3 (Fig. 5a). These pockets have average dynamic volumes of 24, 119, 201.1, 55, and 50 Å^3^, respectively (Fig. S7).

**Fig 5.**
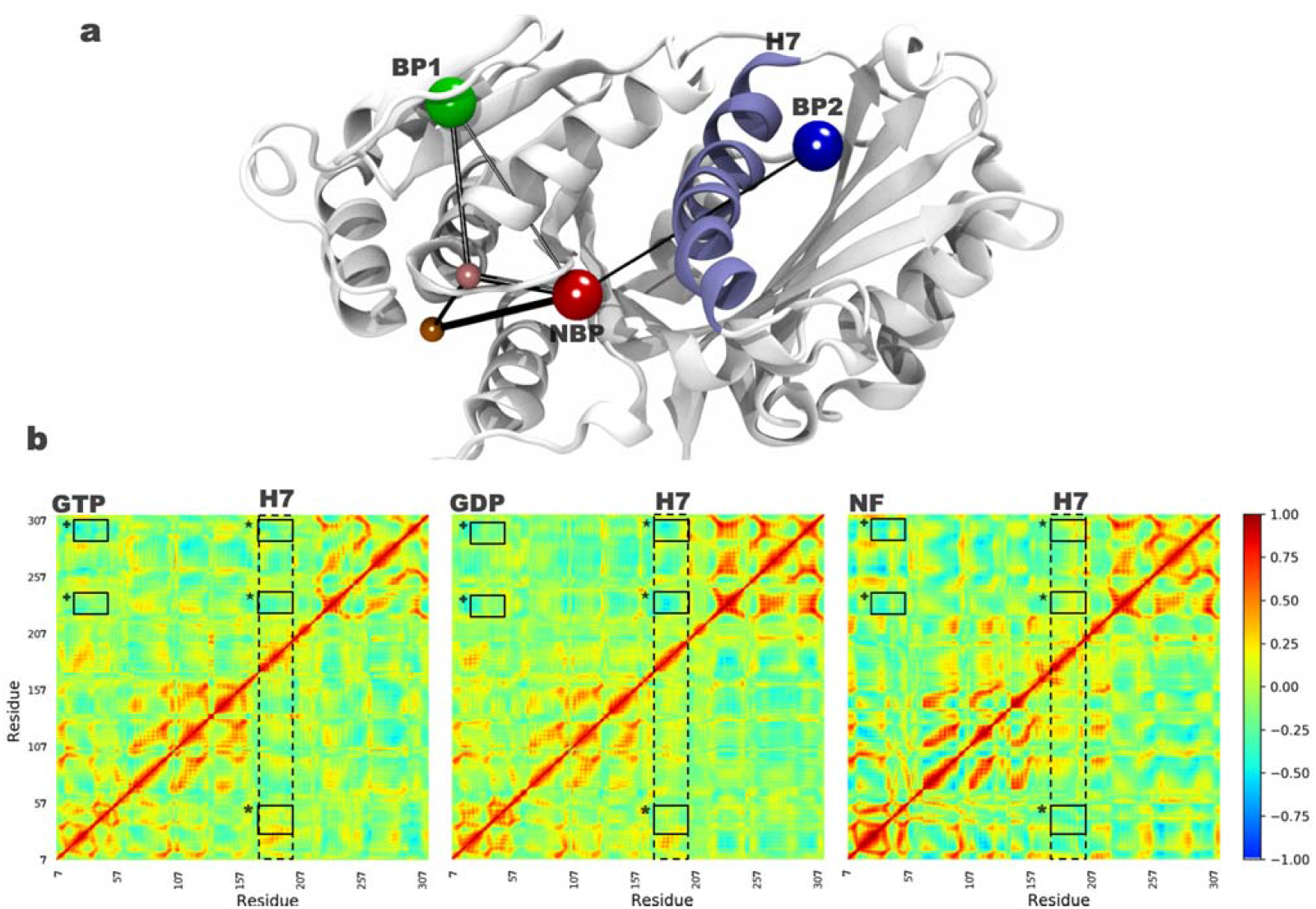
Communication between binding pockets. **a**, Spatial location of 4-hydroxycoumarin binding pocket (BP) 1 (green), 2 (blue), nucleotide-binding pocket (NBP; red) and two additional sub-pockets (salmon and pink) computed for the nucleotide-free FtsZ structure. The pockets are connected via black lines (edges) and exhibit a persistency of at least 24%. Helix H7 is coloured purple. **b**, Dynamic cross-correlation analysis of the time-correlated Cα motions calculated from FtsZ-GTP (left), FtsZ-GDP (centre) and nucleotide-free FtsZ (right) complex simulations. The cross-correlation between helix H7 (dashed box) and BP1 (*), BP2 (D) and between BP1 and BP2 (D) are illustrated proportional to the shading, as indicated by the scale on the right.

The reduced volume of BP1 can be explained by the existence of the nearby nucleotide-binding site, which has a persistence of ∼80%. Furthermore, the high fluctuations of loop T3 in the nucleotide-free state allow the formation of two minor pockets (pID 20 and 14) with a persistence of 45 and 55%. These sub-pockets have the ability to merge with the nucleotide-binding pocket, resulting in a single larger binding pocket (Fig. 5a; Fig. S7). BP2 is situated diagonally across helix H7 on the distal side. Since the conformation of H7 controls the volume of both BP2 and the nucleotide-binding site, it was therefore not surprising that the nucleotide-binding site and BP2 displayed a crosstalk pattern during our simulations. This suggests that a possible communication pathway exists between BP2 and BP1 that is mediated via the nucleotide-binding site. Furthermore, this might also explain the formation and disappearance dependence of BP1 and BP2 on the dynamics of helical bending in the nucleotide-free state. However, in the nucleotide-bound state, the communication between BP2 and BP1 is blocked by the presence of the nucleotide and its strong interactions within its binding site.

### Dynamic cross-correlation between helix H7 and 4-hydroxycoumarin binding pockets

To investigate the relative residue movement between the Cα atoms of BP1, BP2 and helix H7, we generated dynamical cross-correlation (DCC) heat maps (Fig. 5b). The DCC was calculated separately over all 30000 frames for GTP-bound, GDP-bound and nucleotide-free complex. In the nucleotide-bound states, the interactions between BP1 and helix H7 are positively correlated (marked *), whereas the interactions between BP2 and H7 helix (marked □) are anticorrelated. The DCC between BP1 and BP2 (marked □) are also anticorrelated. However, in the nucleotide-free state the DCC between BP1 and helix H7 is anticorrelated, whereas the movement between BP2 and helix H7 is correlated. The interactions between BP1 and BP2 are also partially positively correlated. This is consistent with our pocket crosstalk analysis, which revealed an allosteric communication between BP1 and BP2. Taken together, our analysis confirms that formation of the 4-hydroxycoumarin binding pockets in the nucleotide-free state and its absence in the nucleotide-bound state are a result of cryptic communication in *M. tuberculosis* FtsZ.

## Discussion

The antibacterial effects of coumarin and its analogues have been quantified in various bacterial strains including *S. aureus, E. coli, P. aeruginosa* and *M. tuberculosis H*_*37*_*Rv* [24, 25]. Duggirala and colleagues [26]were able to prove that the 4-hydroxycoumarin inhibited the both the polymerization and GTPase activities of *E. coli* FtsZ, providing a direct link between inhibition of FtsZ and its antibacterial activity. In the absence of structural data for the inhibition of FtsZ by coumarins, modeling attempts have been undertaken to predict the coumarin-binding pocket for *M. tuberculosis* FtsZ [26]. These docking studies using the *E. coli* FtsZ structure as a template predicted that coumarin binds to FtsZ at a site distinct from the GTP binding site, in close proximity to the binding pocket identified in *S. aureus* FtsZ (Fig. S8). This predicted binding pocket is part of the T7 loop of *S. aureus* FtsZ. However, the two new coumarin-binding pockets that we report in this work are not located in the predicted location. The reasons for this discrepancy may be that subtle structural differences between *E. coli, S. aureus* and *M. tuberculosis* FtsZ structures are more significant than anticipated. Moreover, it is worth emphasizing that all previous attempts to model coumarin binding were done on the GTP-bound structure, where the cryptic pockets that we report are not formed.

*S. aureus* FtsZ exists in open and closed conformations [27]. To date, only a single inhibitor-binding pocket has been described by structural methods, namely for *S. aureus* FtsZ [12, 13, 28]. This binding pocket is located in a hydrophobic cleft between the C-terminal subdomain and helix H7 in proximity to the T7 loop and is distinct from the two new pockets identified in our study (Fig. S8). In the *S. aureus* complex (Table S2), the inhibitor PC190723 (or its analogues) is bound to the FtsZ-GDP binary complex in the open conformation. Since there is only one binary FtsZ-GDP complex in the asymmetric unit, it is unclear if this represents a transient inhibitor-binding pocket or if this inhibitor could also bind to nucleotide-free FtsZ. Superimposition of *M. tuberculosis* and *S. aureus* FtsZ closed conformation structure (PDB entry 3WGK) reveals a Cα RMSD of ∼ 0.4 Å (Fig. S9a). This confirms that the coumarin analogues stabilize the closed conformation of *M. tuberculosis* FtsZ, unlike *S. aureus* FtsZ where the inhibitor stabilizes the open conformation. A comparison of *M. tuberculosis* FtsZ with the inhibitor-bound open conformation *S. aureus* FtsZ highlights a significant rotation of the C-terminal domain by ∼27° (Fig. S9b). In addition to this rotation, the H7 helix in *S. aureus* displays a downward shift by about one helical turn (Fig. S9c). It is interesting to note that none of the *S. aureus* structures (open or closed conformations) displayed H7 helical bending or the formation of equivalent binding sites for coumarin analogues. Nevertheless, we do not exclude the possibility that even more binding pockets will be identified in the future in FtsZ.

Bacterial FtsZ also shares homology with mammalian tubulin. Interestingly, certain taxol-related analogues that have been shown to be inactive against various tumour cell lines and do not act on tubulin are still potent inhibitors against *M. tuberculosis* FtsZ with MIC values of 2.5 μM [29]. Comparison of the FtsZ-4-hydroxycoumarin binary complexes with medically relevant tubulin-drug complexes (Fig. 6), namely tubulin in complex with the anticancer agents taxol (PDB entry 1JFF, [30]), colchicine (PDB entry 4O2B, [31]) and vinblastine (PDB entry 5J2T, [32]), highlighted that the two transient 4-hydroxycoumarin binding pockets on FtsZ do not overlap with either colchicine (Fig. 6b) or vinblastine (Fig. 6c). In contrast, one of the 4-hydroxycoumarin binding pockets overlaps with the taxol-binding pocket on tubulin (Fig. 6a), indicating that FtsZ and mammalian tubulin share not only the same ancestor with respect to their overall folds but also some conservation of their inhibitor-binding pockets.

**Fig 6.**
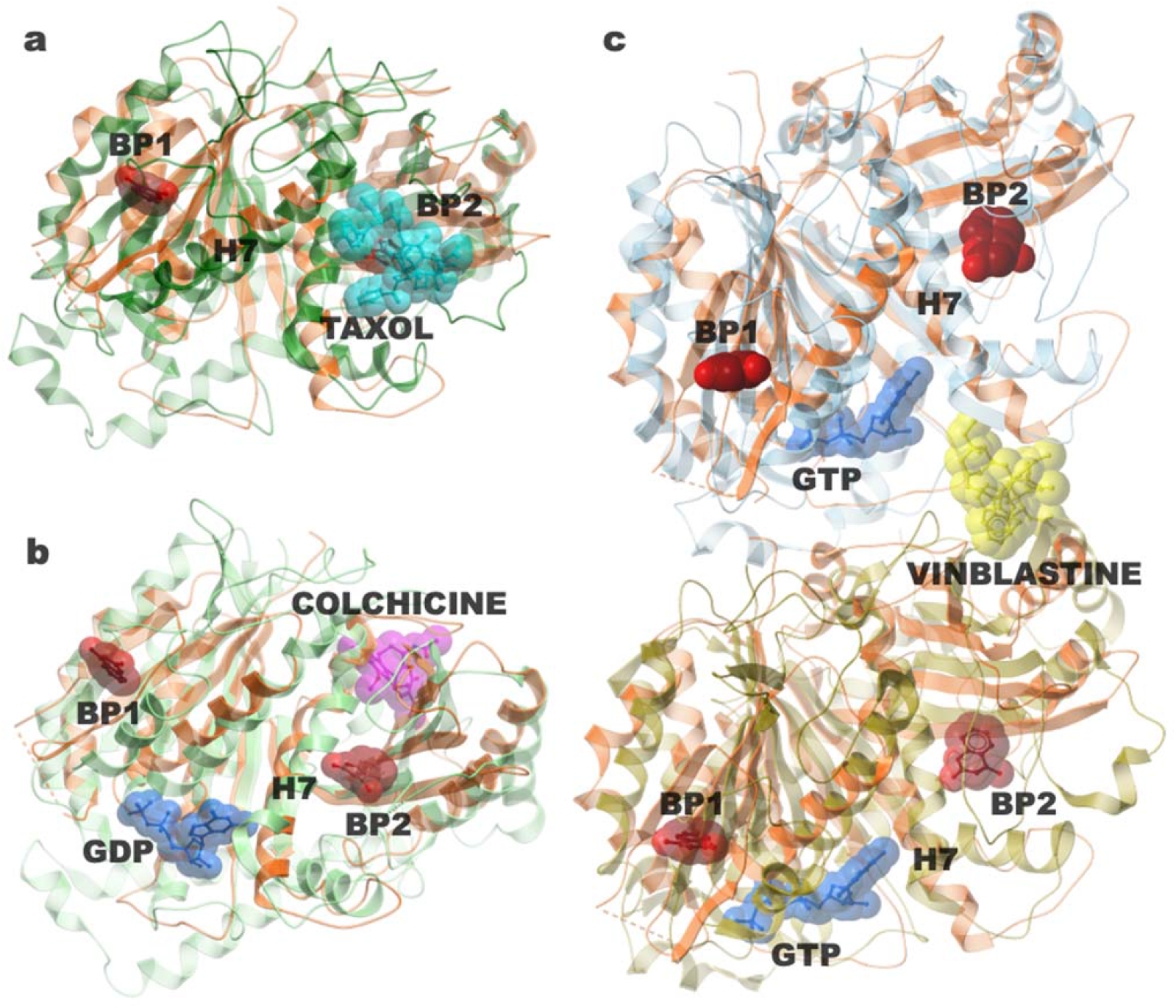
Structural comparison of FtsZ-4-hydroxycoumarin and tubulin-inhibitor complexes. The nucleotide-free FtsZ (orange)-4-hydroxycoumarin (red space fill) complex has been superimposed with clinically relevant (**a)** tubulin-taxol (cyan) complex, (**b)** tubulin-colchicine (pink) complex and (**c)** tubulin-vinblastine (yellow) complexes.

Finally, our results provide novel insights into the structural and dynamic mechanisms into cryptic pocket formation on *M. tuberculosis* FtsZ. We report on the formation and dissolution of two novel pockets on *M. tuberculosis* FtsZ. A combination of crystallographic and computational work has allowed us to understand structural dynamics in a manner that is not evident from static structures alone. The atomistic level details in understanding the cryptic pocket formation not only opens new avenues to study and understand FtsZ function but can also be leveraged to design more potent and selective inhibitors.

## Methods

### Reagents and Constructs

4-Hydroxycoumarin was bought from hit2lead / Chembridge with a purity of > 90%. BL21(D3) pLysS competent cells and Bugbuster protein extraction reagent were purchased from Novagan. LB medium and LB-agar were purchased from MP Biomedicals. TB medium was bought from Fluka. NuPAGE®LDS Sample Buffer (4X), NuPAGE®Sample Reducing Agent (10X), NuPAGE® MES SDS Running Buffer (20X), SeeBlue® Plus2 Pre-stained Protein Standard, SimplyBlue™ SafeStain and NuPAGE®Novex® 4-12% Bis-Tris Protein Gels (1.0 mm 10 well) were purchased from Life Technologies. Tris (hydroxymetthyl) aminomethane, Imidazole, distilled water, MgCl_2_, MgAc_2_, Glycerol, Dimethyl sulfoxide (DMSO), Ethylene glycol tetraacetic acid (EGTA), NaCl, Benzonase Nuclease, Chloramphenicol, Ethylenediaminetetraacetic acid (EDTA), N-(2-acetamido)-2 amino ethanesulphonic acid (ACES), KCl, and Isopropyl-D-1-thiogalactopyranoside (IPTG) were purchased from Sigma Aldrich. Ethanol, NaOH, Sodium citrate dihydrate, and Isopropanol were purchased from Fisher. 5 ml HisTrap FF columns, Superose 12 was bought from GE Healthcare. Ampicillin sodium salt was bought from Cayman Chemical. SnakeSkin^®^ Dialysis Tubing was purchased from Thermo Scientific. PEG4000 was purchased from Melford. Ammonium acetate was purchased from Lab Chemie. 24 well linbro plates and coverslips were bought from Hampton Research.

### Protein expression and purification of FtsZ from *M. tuberculosis*

Full-length FtsZ from *M. tuberculosis* (residues 1-379) was cloned into the pProEx expression plasmid, yielding a PreScission Protease cleavable N-terminal hexahistidine-tagged protein. The plasmid was transformed into BL21(DE3) pLysS. A single colony was mixed into a 20-mL flask containing TB medium, supplemented with 100 µg/ml ampicillin and 34 µg/mL chloramphenicol. The flask was incubated at 37°C for 8 h. The culture was distributed equally into six 2L flasks containing 1 L TB medium, supplemented with 100 µg/mL ampicillin, and cultured overnight at 37°C. The next day, cells were induced by the addition of 0.5 mM IPTG for 24 h at 20°C. Cells were then harvested by centrifugation at 8000 rpm (Avanti® JE centrifuge and JLA-16.250 rotor from Beckman Coulter) for 10 min at 4°C. The supernatant was discarded, and the pellets were stored at −80°C until used. The harvested cells were resuspended in buffer A (50 mM HEPES pH 7.2, 300 mM NaCl, 5% glycerol and 10 mM imidazole) and lysed on ice by sonication, for ten cycles of 30 sec on/off. The lysate was transferred into precooled 250 mL centrifuge bottles and centrifuged at 20,000 rpm (Avanti® JE centrifuge and JLA-16.250 rotor from Beckman Coulter) for 15 min at 4°C.

The resulting supernatant was loaded into a 5 ml His-Trap FF column, equilibrated with 15 mL (1 mL/min) of buffer A (50 mM HEPES pH 7.2, 150 mM NaCl, 5% glycerol and 10 mM imidazole). The column was washed with 100 mL (1 mL/min) washing buffer B (50 mM HEPES pH 7.2, 150 mM NaCl, 5% glycerol and 50 mM imidazole) and the protein was eluted using buffer C (50 mM HEPES pH 7.2, 150 mM NaCl, 5% glycerol and 250 mM imidazole).

The eluted protein was mixed with the appropriate amount of PreScission protease (1 mg of PreScission protease / 50 mg protein), transferred into 10 K SnakeSkin Dialysis Tubing and placed overnight at 4°C into cleavage buffer (50 mM HEPES pH 7.2, 150 mM NaCl, 1 mM EDTA, 0.01% Tween 20 and 1 mM DDT). Any uncleaved His-Tagged protein or PreScission protease was removed by running the protein through a second 5 ml HisTrap FF column. Cleaved FtsZ was dialysed against dialysis buffer (25 mM MES pH 7.2, 0.1 mM EDTA, 10 mM DDT, 50 mM NaCl and 5% glycerol) and concentrated to 18 mg/mL using an Amicon Ultra-15, 10 kDa concentrator, frozen in liquid nitrogen and stored at-80°C.

### Protein crystallisation of native FtsZ and in complex with 4-hydroxycoumarin

Purified FtsZ (18 mg/mL) was crystallised in Linbro plates using the hanging drop method. 1 μl of protein was mixed with a drop of 1 μl of reservoir solution containing 27.5% PEG4000, 0.4 M ammonium acetate and 0.1 M Na citrate pH 5.6 and incubated at 18°C for at least six weeks. For the native *M. tuberculosis* FtsZ structure, crystals were immersed in a cryoprotectant solution composed of reservoir solutions plus 15% glycerol while, for the FtsZ-4-hydroxycoumarin structure, crystals were soaked for 30 min in a cryoprotectant solution composed of liquid mother solution plus 15% DMSO containing 37.5 mM of 4-hydroxyloumarin. Finally, all crystals were flash-frozen in liquid nitrogen and stored until used.

### Data collection, structure determination and refinement

Diffraction data for native *M. tuberculosis* FtsZ and the two coumarin analogue complexes were collected at Diamond Light Source, Oxfordshire, at beamline i04. Data were processed using iMosflm [33] and scaled using SCALA [34] to resolutions as shown in Table 1. The structures of native FtsZ and the FtsZ-4-coumarin complexes were solved in space group P_65_ by molecular replacement (PHASER MR in CCP4 suite [35] using the previously published *M. tuberculosis* FtsZ structure (PDB ID: 1RQ7;[36]) as a search model. All structures were improved using Coot and refined with PHENIX [37]. The calculation of R_free_ used 5% of the data. Crystallographic and refinement statistics are shown in Table 1. Native FtsZ contains residues Asn6 to Thr63 and Gly69 to Phe312 in chain A, whereas chain B contains Asn6 to Gly59 and Ala70 to Asp313.

### Molecular Dynamics (MD) Simulations

The starting structures for molecular dynamics simulations were prepared after the final crystallographic refinement of the resolved coordinates. Three separate systems GTP, GDP and nucleotide-free (NF) were prepared as follows: While preparing the NF state, the bound coumarin ligands in binding pockets 1 and 2 were removed. The protonation state of the complexes was determined using the software PDB2PQR [38]. This was followed by the systems being set up using xleap [39] employing the ff14SB forcefield [40]. The parameters for GTP and GDP were downloaded from the Amber parameter database [41]. The protonated complexes were solvated using TIP3P water [42] and the edge of the box was set to at least 10 Å form the closest solute atom. The system was neutralised using counter ions. The protocol was identical for each system. Each system was minimized and equilibrated under NPT conditions for 5 ns at 1 atm. The temperature was increased gradually to 298 K using a timestep of 4 fs, rigid bonds, a cut-off of 9 Å and particle mesh Ewalds summation switched on for long-range electrostatics. The protein was fixed during equilibration, while the ions and solvent molecules were allowed to move. The heavy atoms of the protein and co-factors were constrained by a spring constant set at 1 kcal mol^−1^ Å^−2^. The production simulations were run using ACEMD molecular dynamics software [43] in the NVT ensemble using a Langevin thermostat with a damping of 0.1 ps^−1^ and hydrogen mass repartitioning scheme to achieve time steps of 4 fs. Two sets of simulations were run for each system for 3 μs (30000 frames) starting from different velocities to improve statistics. VMD [44] was used to visualise the trajectories and the figures were made using ICM-Pro software (www.molsoft.com).

### Clustering

Trajectories from all three simulations (sampling 9 μs) were pooled together before clustering was carried out. All frames were aligned and grouped based on the RMSD metric, such that similar ones are in the same cluster while the dissimilar structures are in other clusters. Clustering was done via the *k-medoids* algorithm [14] with a smart initialisation method implemented in the BiKi Lifesciences software suite (www.bikitechnologies.com). The representative conformation called medoids is based on the most centrally located conformation in the cluster and is less sensitive to outliers in comparison with other methods. The size of the circle is proportional to the size of the cluster and the edges denote the number of inter-conversions between them. The Fruchterman-Reignold layout was used to generate the cluster graph.

### Helical bending

The dynamics of helix H7 was analysed using Bendix software [45] implemented in the Visual Molecular Dynamics package [44]. The Bendix algorithm is based on calculating a window of four residues to give local helix axes that are joined by a spline [46]. The helical bending is plotted as a heatmap, which is colour coded according to the local angle magnitude, highlighting non-linear helix behaviour. The maximal and local angles were plotted over the course of each simulation set.

### Pocket crosstalk

Spatio-temporal evolution of the binding pockets was investigated using Pocketron [23] implemented in the BiKi Lifesciences software suite (www.bikitechnologies.com). The algorithm identifies pockets and tracks their evolution over time along an MD trajectory. In brief, residue exchange between adjacent pockets, termed as pocket crosstalk, was monitored. This is another means of identifying communication between different parts of the protein [21, 22]. Alignment-independent identification of pockets on the protein surface was carried out, followed by quantification and visualisation of the volume and surface area of all the pockets found in the protein, for each conformation present in the MD trajectory. Each pocket is then assigned a unique pocket identifier (pID). Communication between pockets, describing the merging and splitting is plotted. More specifically, a “merge” event occurs when atoms belonging to the same pocket in the current frame have belonged to separate pockets in a previous frame. Similarly, a “split” event occurs when atoms of a single pocket separate into two or more distinct pockets. The algorithm translates the merging and splitting frequency matrix into a 3D network graph, where nodes represent geometric center of the pockets, and the edges indicate communication between two pockets. The thickness of the edge is directly proportional to the frequency of the merging and splitting events between two connected pockets. Finally, the signal transmission network between pockets across the protein surface is illustrated as interconnected pocket motions. The entire 3 μs nucleotide-free trajectory with a stride of 2 (>15000 frames) was used to analyse pocket crosstalk for each system. The minimum persistence time for each pocket was fixed at 24%.

### Dynamic cross-correlation maps

The systems were simplified by representing each residue as a single Cα node. All 30000 conformations pooled for each system was used to construct a local contact matrix. Two nodes (excluding neighbouring nodes) are in contact if they are within a distance of 6.5 Å. The connectivity between two nodes is termed an edge. The dynamic cross-correlation maps (DCCM) were calculated using the program MD-TASK program [47]. The generated dynamic cross-correlational map has the ability to identify highly correlated or anticorrelated nodes. However, to compute communication pathways, it is useful to construct a matrix (C) where small values indicate highly correlated or anticorrelated motions. The edges between the nodes can be weighted on how often the interactions exist. This can be functionalised by w_ij_ = −log(|C_ij_|); where w_*ij*_ can be thought of as a distance in the functionalised correlation space between node-node pairs *i* and *j*.

## Data Availability

The structures of native FtsZ in the presence of GDP (PDB ID 6YM1) or GTPyS (PDB ID 6YM9) as well as FtsZ-GDP-coumarin (PDB ID 6Y1U) and FtsZ-GTPyS-coumarin (PDB ID 6Y1V) ternary complexes can be downloaded from the Protein Data Bank. Simulation trajectories and all other data supporting the findings of this study are available from the corresponding author upon reasonable request.

## Reporting summary

Further information is available in the Nature Research Reporting Summary linked to this article

## Acknowledgements

We are grateful for PharmAlliance for initial seed funding and the Faculty of Pharmacy, King Abdul-Aziz University for Aisha Alnami’s PhD studentship. We thank Diamond Light Source for providing access to beamlines I04 and I24, which contributed to the results.

## Author contributions

The manuscript was written through contributions of all authors. All authors have approved the final version of the manuscript. This publication is part of the doctoral thesis of Aisha Alnami.

## Additional information

**Supplementary information** is available for this paper.

**Correspondence and requests for materials** should be addressed to F.K.

