## Supplementary Information for "Conformational flexibility Of A Highly Conserved Helix Controls Cryptic Pocket Formation In FtsZ"

### Summary

Table S1: Data collection, structure determination and refinement statistics for native and coumarin-bound FtsZ in the absence and presence of GDP 2

Table S2: Data collection, structure determination and refinement statistics for native and coumarin-bound FtsZ in the absence and presence of GTPγS 3

Table S3: Published *M. tuberculosis* FtsZ structures deposited in the PDB 4

Table S4: Published *S. aureus* FtsZ-inhibitor complexes 5

Fig S1: Multiple sequence alignment of FtsZ sequences 6

Fig S2: 2FoFc maps of BP1 and BP2 7

Fig S3: Water molecules around BP1 and BP2 8

Fig S4: Clustering analysis of the MD simulation trajectories 9

Fig S5: Details of the representative structures from clusters 10

Fig S6: Overlay of the representative structures from clusters 11

Fig S7: Dynamic volume of the five binding pockets 12

Fig S8: Comparison of binding pockets identified in *S. aureus* ternary FtsZ-GDP-PC190723 and *M. tuberculosis* FtsZ-4-hydroxycoumarin complexes 13

**Fig S9**. Comparison of the closed and open conformations of *S. aureus* FtsZ with *M. tuberculosis* FtsZ-4-hydroxycoumarin complex 14

**Table S1** Data collection, structure determination and refinement statistics for native *M. tuberculosis* FtsZ in the absence and presence of GDP and the binary FtsZ-4-hydroxycoumarin complex. Statistics for the highest-resolution shell are shown in parentheses.

| Parameters | Native FtsZ GDP-bound & nucleotide-free | FtsZ-GDP-4-hydroxycoumarin complex |
|---|---|---|
| **PDB ID** | 6YM1 | 6Y1U |
| **Beamline** | I24 | I24 |
| **Resolution range** | 76.68 - 1.70 (1.761 - 1.70) | 70.07 - 1.68 (1.74 - 1.68) |
| **Space group** | $P6_5$ | $P6_5$ |
| **Unit cell parameters** | 88.5, 88.5, 178.1, 90, 90, 120 | 88.1, 88.1, 176.8, 90, 90, 120 |
| **Molecules in AU** | 2, GDP-bound & nucleotide-free | 2, GDP-bound & nucleotide-free |
| **Total reflections** | 690527 (197096) | 1021303 (129960) |
| **Unique reflections** | 86493 (8594) | 88000 (8733) |
| **Multiplicity** | 8.0 (7.7) | 11.6 (10.2) |
| **Completeness (%)** | 100.0 (100.0) | 99.8 (99.5) |
| **Mean I/sigma(I)** | 9.5(1.8) | 17.5 (2) |
| **Wilson B-factor** | 30.1 | 25.0 |
| **R-merge** | 0.093 (0.991) | 0.077 (1.184) |
| **R-meas** | 0.100 (1.062)) | 0.080 (1.248) |
| **CC1/2** | 0.998 (0.668) | 0.999 (0.479) |
| **Reflections used in refinement** | 86199 (8468) | 87997 (8732) |
| **Reflections used for R-free** | 4299 (442) | 4365 (491) |
| **R-work [%]** | 18.6 (30.6) | 17.7 (30.0) |
| **R-free [%]** | 21.7 (33.6) | 20.3 (32.5) |
| **No of non-hydrogen atoms** | 4669 | 4853 |
| **Macromolecules** | 4296 | 4347 |
| **Ligands** | 28 | 132 |
| **Solvent** | 598 | 374 |
| **Protein residues** | 598 | 600 |
| **RMS (bonds)** | 0.013 | 0.012 |
| **RMS (angles)** | 1.51 | 1.29 |
| **Ramachandran favoured (%)** | 99.5 | 98.7 |
| **Ramachandran allowed (%)** | 0.5 | 1.0 |
| **Ramachandran outliers (%)** | 0.0 | 0.3 |
| **Rotamer outliers (%)** | 1.6 | 1.1 |
| **Clashscore** | 4.6 | 6.0 |
| **Average B-factor** | 36.3 | 30.7 |
| **Macromolecules** | 35.8 | 29.5 |
| **Ligands** | 24.6 | 44.6 |
| **Solvent** | 43.6 | 40.1 |

**Table S2:** Data collection, structure determination and refinement statistics for native *M. tuberculosis* FtsZ in the absence and presence of GTPγS and the binary FtsZ-4-hydroxycoumarin complex. Statistics for the highest-resolution shell are shown in parentheses

| Parameters | Native FtsZ GTPyS-bound & nucleotide-free | FtsZ-4-hydroxycoumarin complex |
|---|---|---|
| **PDB ID** | 6YM9 | 6Y1V |
| **Beamline** | I04 | I04 |
| **Resolution range** | 181.33 - 2.03 (2.03 - 2.03) | 180.30 - 2.53 (2.4 - 2.4) |
| **Space group** | $P6_5$ | $P6_5$ |
| **Unit cell parameters** | 89.1, 89.1, 181.3, 90, 90,120 | 88.5, 88.5, 180.3, 90, 90, 120 |
| **Molecules in AU** | 2, GTPγS-bound & nucleotide-free | 2, GTPγS-bound & nucleotide-free |
| **Total reflections** | 297387 (43170) | 134609 (18839) |
| **Unique reflections** | 51913 (5104) | 31201 (3083) |
| **Multiplicity** | 5.7 (5.8) | 4.3 (4.1) |
| **Completeness (%)** | 99.0 (98.3) | 100 (99.9) |
| **Mean I/sigma(I)** | 9.3 (1.5) | 6.8 (1.3) |
| **Wilson B-factor** | 34.1 | 39.8 |
| **R-merge** | 0.095 (1.101) | 0.154 (1.040) |
| **R-meas** | 0.105 (1.211) | 0.175 (1.193) |
| **CC1/2** | 0.998 (0.565) | 0.992 (0.520) |
| **Reflections used in refinement** | 51848 (5095) | 31183 (3080) |
| **Reflections used for R-free** | 2745 (278) | 1633 (179) |
| **R-work [%]** | 18.0 (24.0) | 17.6 (24.2) |
| **R-free [%]** | 22.0 (27.6) | 23.1 (33.5) |
| **No of non-hydrogen atoms** | 4573 | 4479 |
| **Macromolecules** | 4260 | 4237 |
| **Ligands** | 32 | 92 |
| **Solvent** | 464 | 281 |
| **Protein residues** | 603 | 602 |
| **RMS (bonds)** | 0.012 | 0.013 |
| **RMS (angles)** | 1.37 | 1.25 |
| **Ramachandran favoured (%)** | 99.0 | 98.0 |
| **Ramachandran allowed (%)** | 0.8 | 1.9 |
| **Ramachandran outliers (%)** | 0.2 | 0.2 |
| **Rotamer outliers (%)** | 0.7 | 2.4 |
| **Clashscore** | 3.5 | 4.4 |
| **Average B-factor** | 37.8 | 42.2 |
| **Macromolecules** | 37.6 | 42.1 |
| **Ligands** | 30.3 | 46.7 |
| **Solvent** | 42.2 | 43.9 |

**Table S3:** Published *M. tuberculosis* FtsZ structures. [2]The glycerol is not located in the nucleotide-binding pocket.

| PDB ID | Space group | Molecules in AU | Resolution / $R_{free}$ | Molecule A | Molecule B | References |
|---|---|---|---|---|---|---|
| **1rq2** | $P6_5$ | 2 | 1.86 / 22.2 | [1]Citrate | No nucleotide | [1] |
| **1rlu** | $P6_5$ | 2 | 2.08 / 22.4 | GTPγS | [2]Glycerol | [1] |
| **1rq7** | $P6_5$ | 2 | 2.60 / 24.2 | GDP | No nucleotide | [1] |
| **5ZUE** | $P6_522$ | 1 | 2.70 / 25.3 | GTP | | [2] |
| **4KWE** | $P6_422$ | 3 | 2.91 / 26.7 | GDP | | [3] |

[1]The citrate molecule is located in the nucleotide-binding pocket, near the phosphate positions.

**Table S4:** Published *S. aureus* FtsZ-inhibitor complexes. All complexes contain one molecule of the FtsZ-GDP-inhibitor ternary complex in the asymmetric unit.

| PDB Entry | Molecules in AU | Resolution [Å] | Nucleotide-bound | Comments | References |
|---|---|---|---|---|---|
| **3vob** | 1 | 2.7 | GDP, no $Mg^{2+}$ | Same inhibitor as **4dxd** | [4] |
| **4dxd** | 1 | 2.0 | GDP, no $Mg^{2+}$ | Same inhibitor as **3vob** | [5] |
| **5xdt** | 1 | 1.3 | GDP, no $Mg^{2+}$ | Inhibitor analogue | [6] |
| **5xdu** | 1 | 2.0 | GDP, no $Mg^{2+}$ | Inhibitor analogue | [6] |
| **5xdv** | 1 | 1.7 | GDP, no $Mg^{2+}$ | Inhibitor analogue | [6] |

**Fig S1**

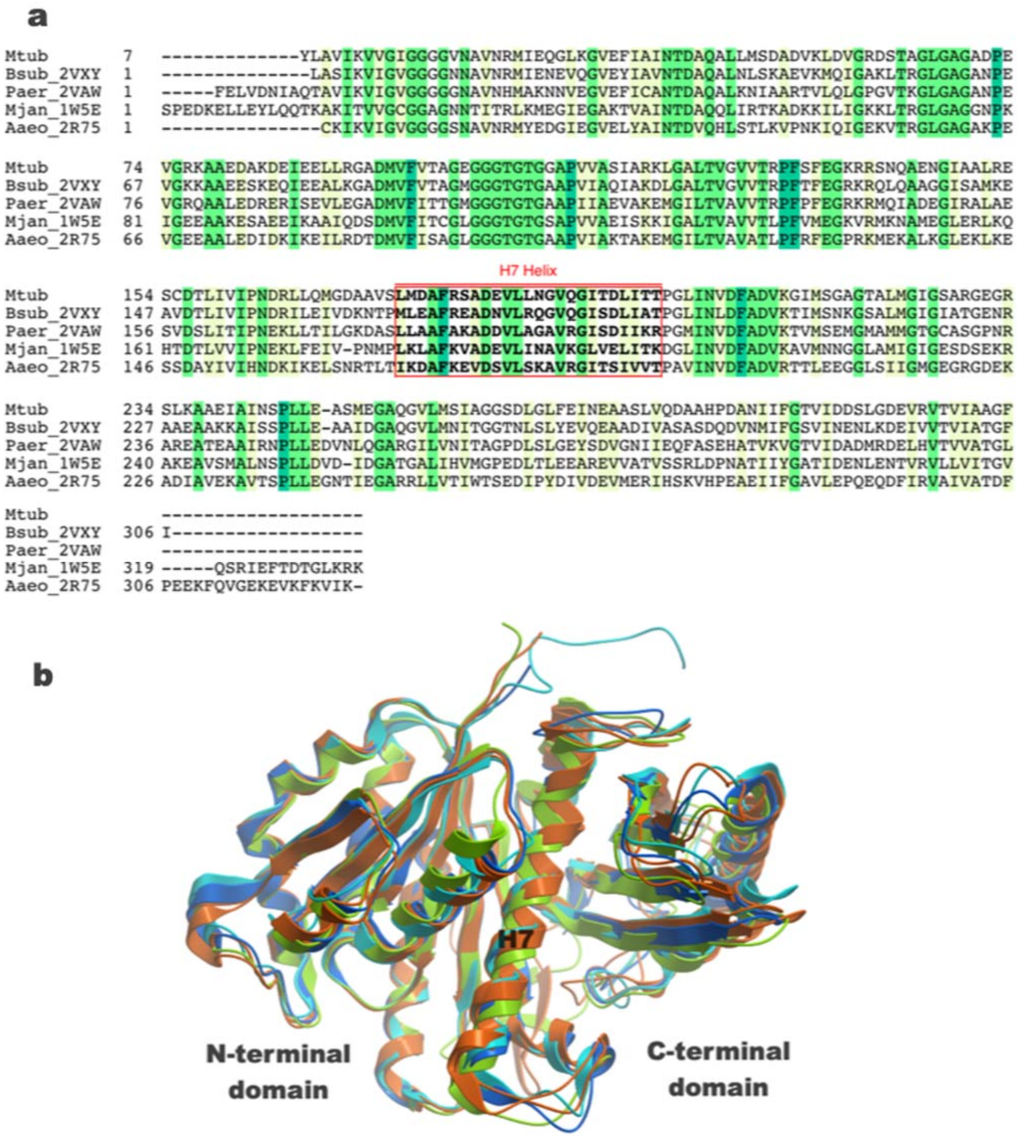


**Fig S1.** (a) Multiple sequence alignment of FtsZ sequences extracted from bacterial crystal structures. Sequence conservation between different bacterial species range between 45-65%. The position of helix H7 labelled (red box) in *M. tuberculosis* (current structure), *B. subtilis* (PDB id 2VXY), *P. aeruginosa* (PDB id 2VAW), *M. jannaschii* (PDB entry 1W5E) and *A. aeolicus* (PDB entry 2R75). (b) Superimposition of the bacterial FtsZ structures. The structural conservation is >0.69 Å.

**Fig S2**

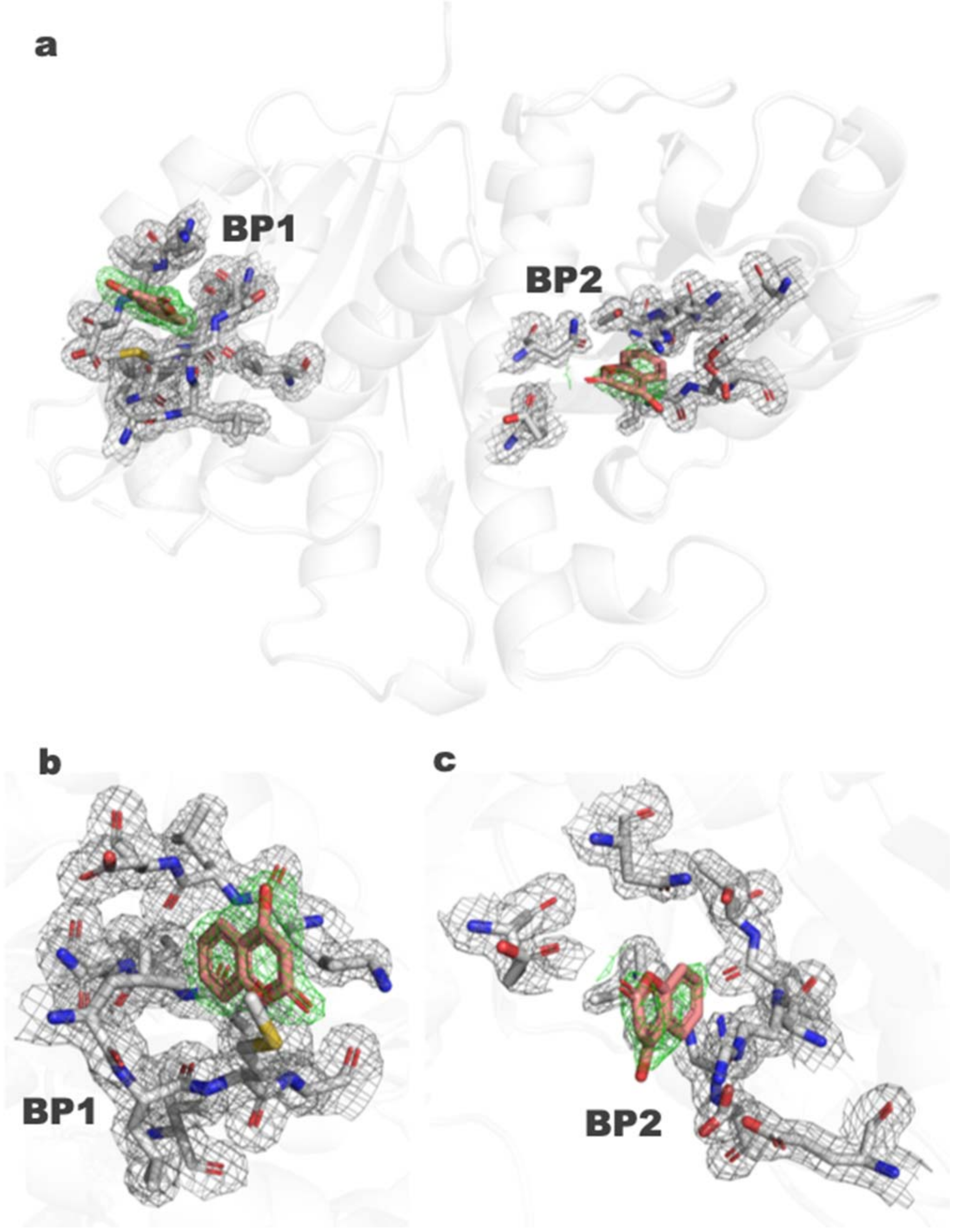


**Fig S2.** 2FoFc maps (1σ) highlighting the electron density around binding pockets 1 and 2 on the nucleotide-Free FtsZ. The electron density around 4-hydroxycoumarin is coloured in green mesh, while the protein side chain electron density is coloured in grey.

**Fig S3**

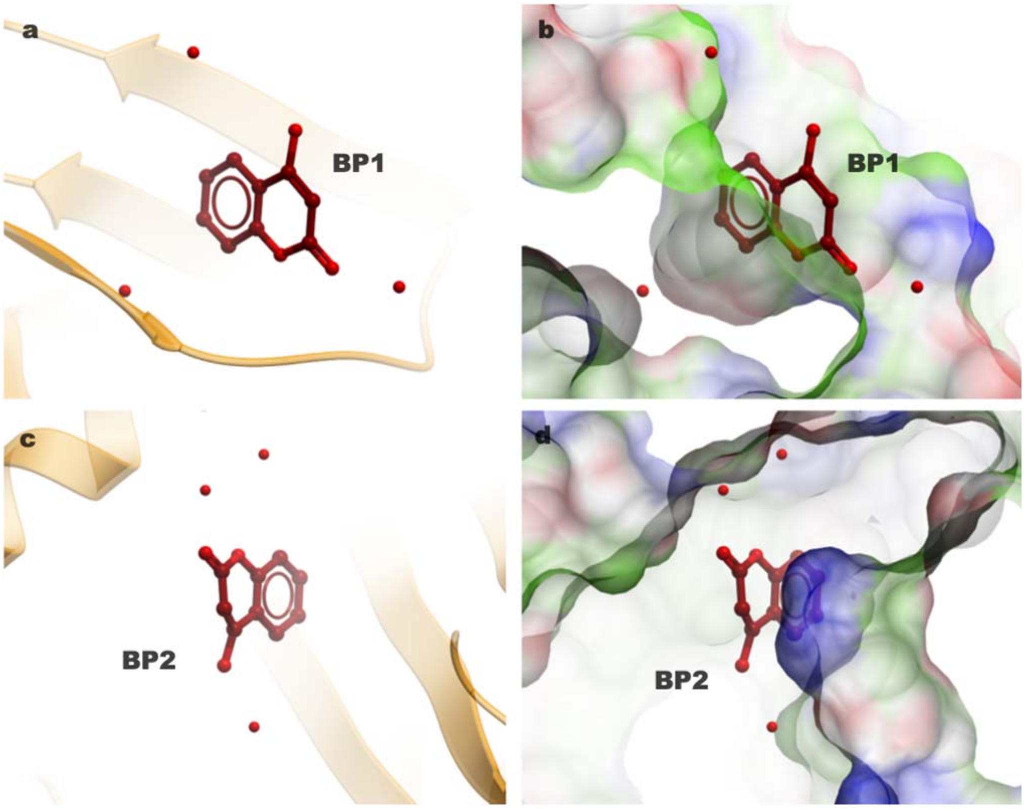


**Fig S3.** Water molecules (red balls) in the vicinity of binding pockets (a,b) 1 and (c,d) 2.

**Fig S4**

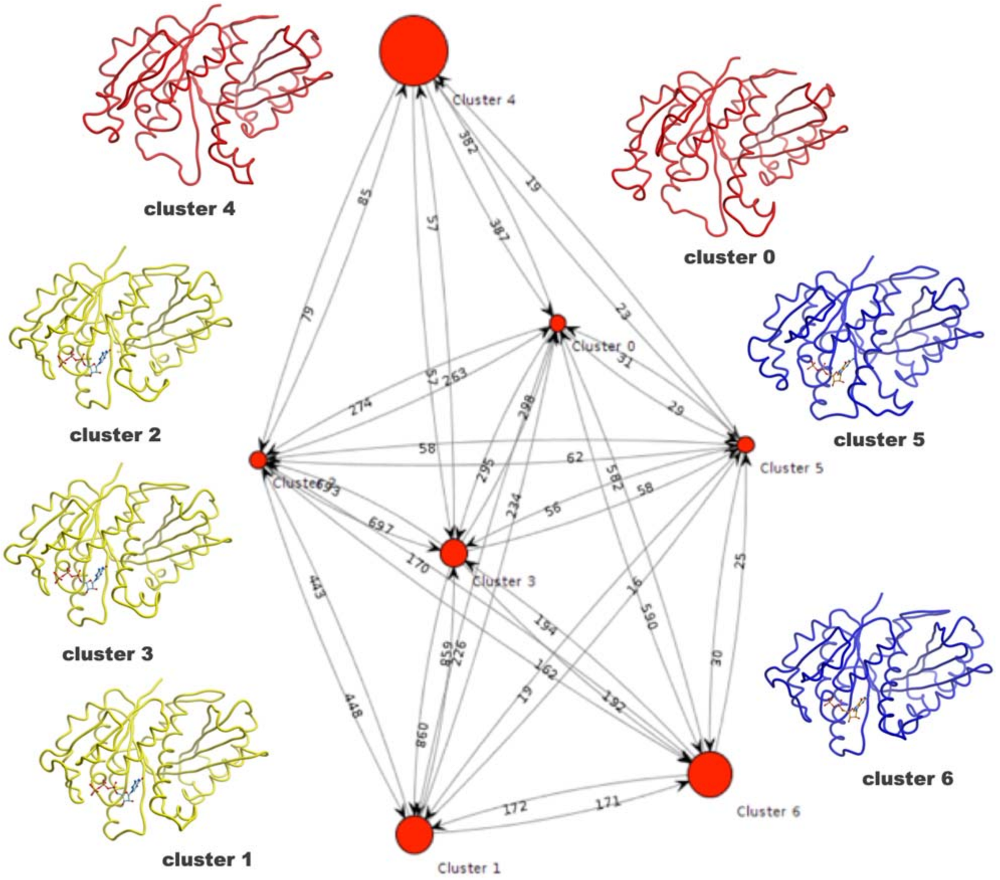


**Fig S4.** Clustering analysis of the MD simulation trajectories via k-medoids algorithm. Circle dimensions represent the cluster size. The number of transitions between each cluster is reported as an edge. The representative structures belonging to their respective cluster are illustrated as protein worms. The representative structures belong to nucleotide-free (red), FtsZ-GDP (blue) and FtsZ-GTP (yellow) trajectories. In total seven clusters were identified, each based on the conformation of the medoid. The size of the circle is proportional to the size of the cluster and the edges denote the number of transitions between them. Cluster 4 represents the nucleotide-free state, Cluster 1 the GTP-bound protein, while Cluster 6 was the GDP-bound state. There is no direct edge connecting the nucleotide-free and the nucleotide-bound states. To go from a nucleotide-bound to a nucleotide-free state, the protein has to take one of the four different routes that pass via one of the four intermediate conformations (clusters 0, 2, 3 or 5). Route 6-0-4 is the most favoured as indicated by the number of edges connecting these cluster centres.

**Fig S5**

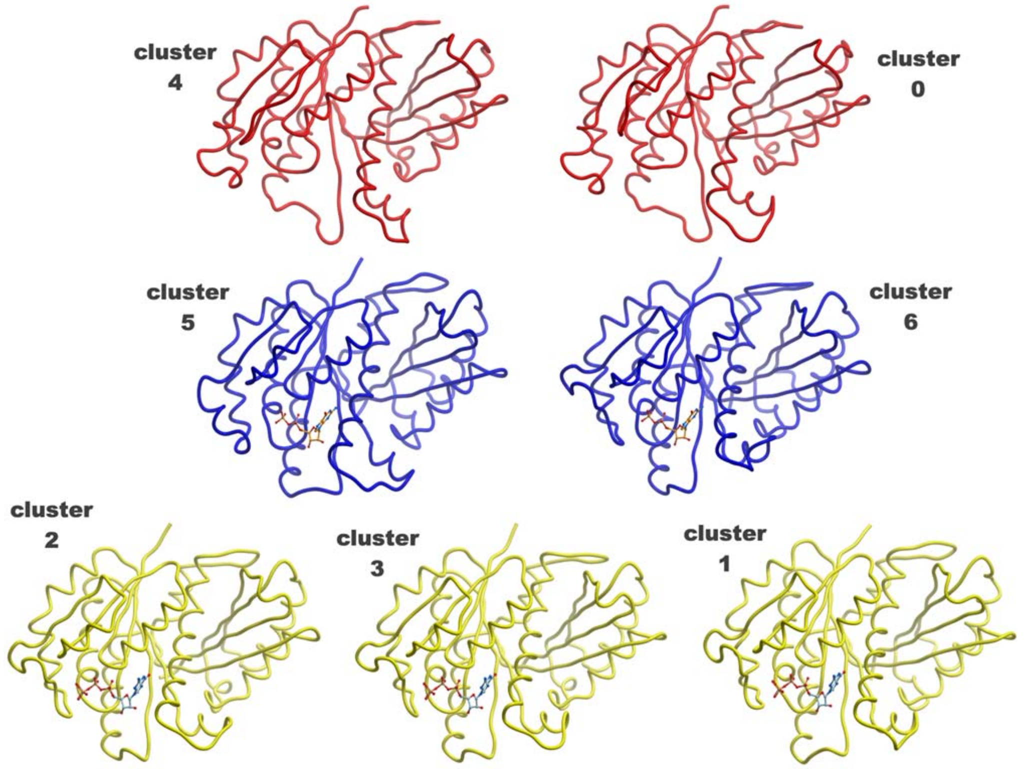


**Fig S5.** Details of the representative structures of their respective cluster illustrated as protein worms. The representative structures belong to nucleotide-free (red), FtsZ-GDP (blue) and FtsZ-GTP (yellow) trajectories.

**Fig S6**

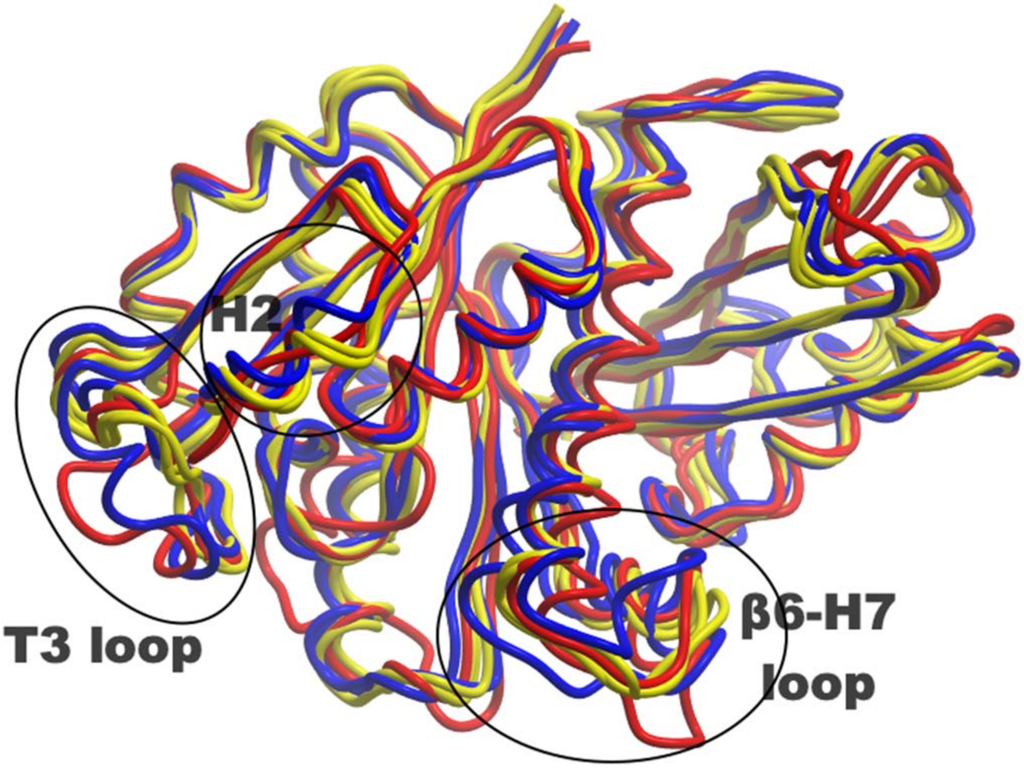


**Fig S6.** Overlay of the representative structures from clustering analysis. The flexible regions are encircled and labelled. The representative structures belong to nucleotide-free (red), FtsZ-GDP (blue) and FtsZ-GTP (yellow) trajectories.

**Fig S7**

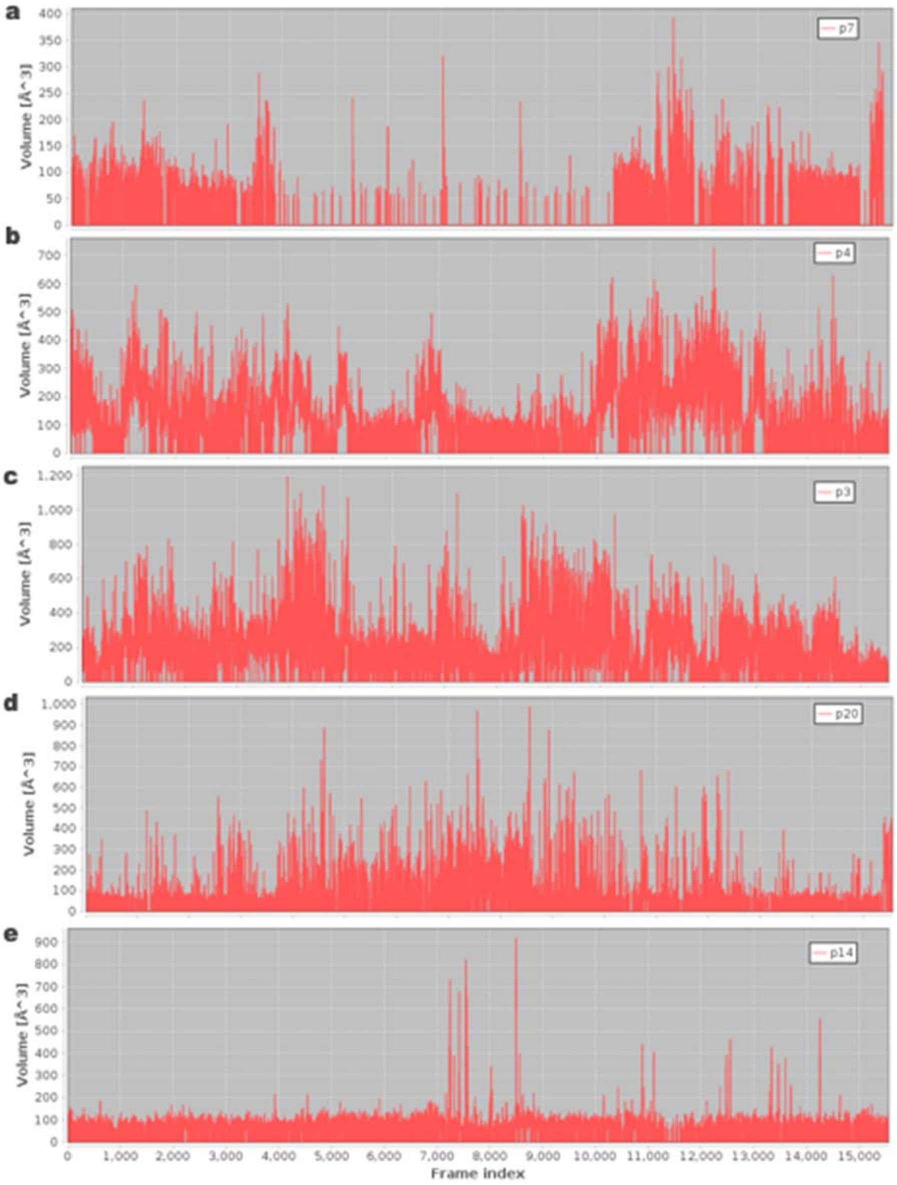


**Fig S7.** Dynamic volume of the five binding pockets calculated from the simulated nucleotide-free trajectory. p7 represents binding pocket 1, p4 represents binding pocket 2 and p3 represents the nucleotide binding pocket. The volume was calculated from 3 μs sampling with a stride of 2, yielding 15000 frames.

**Fig S8**

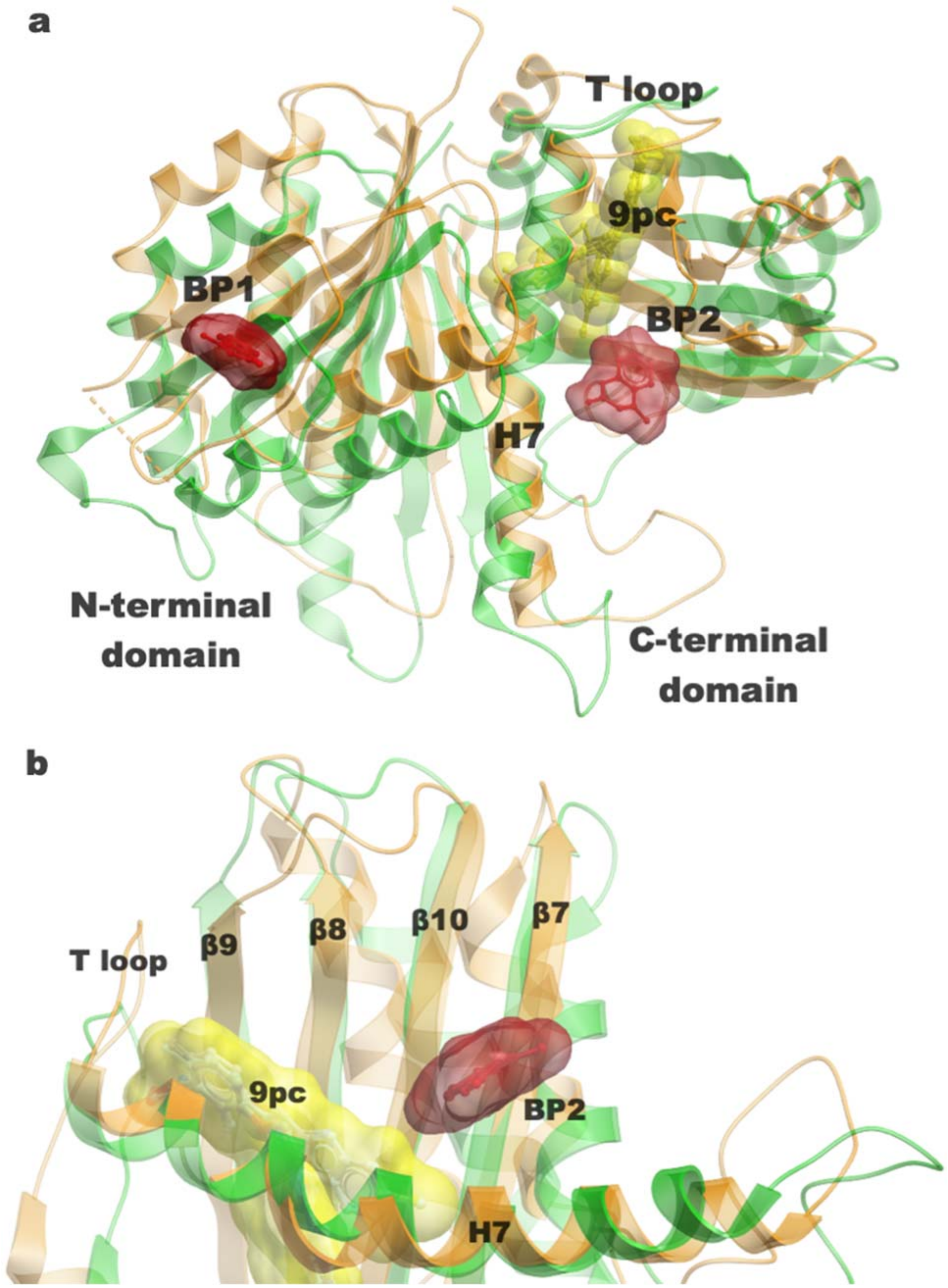


**Fig S8**. Comparison of inhibitor-binding pockets identified in *S. aureus* ternary FtsZ-GDP-PC190723 complex and *M. tuberculosis* FtsZ-4-hydroxycoumarin complex. (a) Inhibitors bound in *S. aureus* (green) and *M. tuberculosis* (orange) FtsZ are illustrated as surface representation. (b) Close-up view of binding pocket 2. PC190723 (9pc) bound to *S. aureus* is illustrated in yellow surface, while 4-hydroxycoumarin is coloured in red.

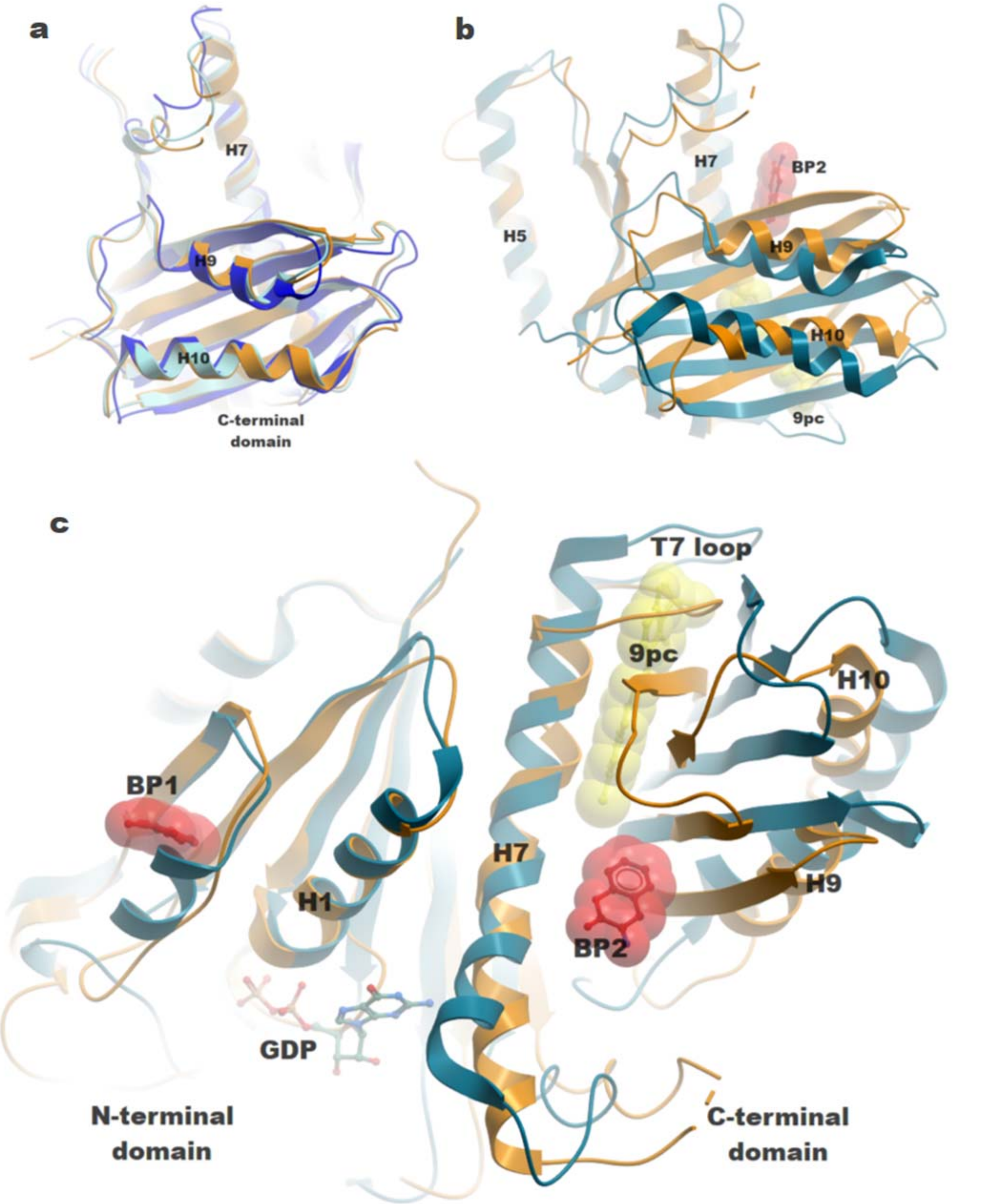


**Fig S9**. Comparison of the closed and open conformations of *S. aureus* FtsZ with the *M. tuberculosis* FtsZ-4-hydroxycoumarin complex. (a) *M. tuberculosis* FtsZ (nucleotide-bound, cyan and nucleotide-free, orange) superimposed on the *S. aureus* FtsZ closed conformation (PDB entry 3WGK; blue) with a Cα RMSD of ~0.4 Å. (b) Superimposition of the *M. tuberculosis* FtsZ-4-hydroxycoumarin complex (orange) on the *S. aureus* ternary FtsZ-GDP-PC190723 inhibitor complex (teal) shows a rotation of the C-terminal domain by ~27°. (c) Overview of the *M. tuberculosis* and *S. aureus* FtsZ complexes highlighting the slide of the H7 helix and T7 loop in the open conformation in *S. aureus* to form the 9pc binding site.

**Movie S1:** Cryptic pocket formation (FtsZ.mov). The short 22 sec movie illustrates the conformational dynamics of the highly conserved helix H7 (coloured in red). Helix H7 is flexible and prone to helical bending. When helix H7 is straight (as observed in the nucleotide-bound states), helix H2 occludes BP1 (shaded in green) and prevents the ligand from binding. In the nucleotide-free state, helix H7 is bent towards the C-terminal domain, away from the nucleotide-binding site. This suggested that helix H2, which forms the boundary of binding pocket 1 (BP1), behaves like a constrained spring that is wound when helix H7 is straight (nucleotide-bound states) and relaxed into the β2'-strand when helix H7 moves away towards the C-terminal domain in the nucleotide-free state. The linked inter-conversion of helix H2 and strand β2', concomitant with helix H7, bending results in the formation or disappearance of BP1.
